# Enhanced long-term memory and increased mushroom body plasticity in *Heliconius* butterflies

**DOI:** 10.1101/2023.07.11.548546

**Authors:** Fletcher J. Young, Amaia Alcalde, Lina Melo-Flórez, Antoine Couto, Jessica Foley, Monica Monllor, W. Owen McMillan, Stephen H. Montgomery

**Author notes:** Corresponding authors: Fletcher J. Young, Stephen H. Montgomery, **Email:** FJY; SHM. **Author Contributions:** SHM designed research; FJY, AA, LM, AC, JF, MM performed research under the supervision of SHM and WOM; FJY analysed data; SHM and WOM provided resources; FJY and SHM wrote the paper with input from all co-authors. **Competing interest statement:** The authors declare no competing interest.

## Abstract

As highly labile structures on both individual and evolutionary time-scales, the mushroom bodies, a key site of learning and memory in insects, are an excellent model for investigating the evolution of cognitive variation. We explored the behavioural consequences of mushroom body expansion in *Heliconius* butterflies, which possess greatly expanded mushroom bodies relative to their closest outgroups. We conducted long-term visual memory assays across three *Heliconius* and three other Heliconiini species using trained food-colour associations. We confirm robust differences between clades, with *Heliconius* exhibiting greater fidelity to the trained colour after 8 days without reinforcement compared to other Heliconiini, with further evidence of stable preferences at 13 days. We extended this analysis to consider the plastic response of the mushroom body calyces over this time period, measuring the volume of the mushroom body calyx, and the number of neurons and synapses it contains. We find substantial post-eclosion expansion and synaptic pruning in calyx of *Heliconius erato*, but not in *Dryas iulia*. In *Heliconius erato*, visual associative learning experience specifically is associated with a greater retention of calyceal synapses. At an individual level, fidelity to the trained colour in *Heliconius erato* was also positively correlated with synapse number. These results point to an enhanced visual long-term memory across *Heliconius*, facilitated not only by phylogenetic expansion of the mushroom body, but also changes in its developmental response to learning experience. The co-evolution of mushroom body expansion, plasticity and specific behaviours provides an important case study in the evolution of cognition.

**Significance Statement:** How are cognitive differences between species supported by evolutionary changes in the brain? We investigated this question using *Heliconius* butterflies which have expanded mushroom bodies, a region of the insect brain involved in learning and memory. We show that *Heliconius* have more stable visual long-term memories and exhibit more substantial age- and experience-related plasticity than a closely related genus with smaller mushroom bodies. Recall accuracy was also predicted by synapse number in *Heliconius erato*, but not *Dryas iulia*, suggesting functional importance. These results suggest that increases in the size of specific brain regions and changes in their plastic response to experience may co-evolve to shape the evolution of cognition.

## Introduction

Animals vary markedly in cognitive ability, both within and between species, yet the neural traits determining these differences are only partly understood, particularly in an evolutionary context (1). A major strand of research has focused on linking measures of “intelligence” to brain size (2–4). Indeed, several studies have found correlations between brain size and certain cognitive abilities (5–7). However, attempts to link brain size to cognition, defined broadly as “the mechanisms by which animals acquire, process, store, and act on information from the environment” (8), have been also critiqued, particularly in an interspecific context (9–11). Whole brain size can ignore variation in brain composition and connectivity (12–15) and non-linear scaling relationships between brains and behaviour (16). Comparative behavioural studies may also fail to consider important ecological differences between species, casting doubt over the biological relevance of the assessed cognitive tasks (10, 17).

Many of these issues can be avoided by conducting studies within species (18–20), or among closely related species (21), focusing on linking specific neural traits and cognitive abilities (10). A classic example is the enlargement of the hippocampus in food-storing birds and its role in spatial memory (22). In insects, the mushroom bodies have similarly been a focus of investigation as a higher-order integrative centre for learning and memory (23, 24). The mushroom bodies have undergone expansion in several insect lineages including Hymenoptera (13), cockroaches (25), herbivorous scarab beetles (26) and *Heliconius* butterflies (27–29). However, the behavioural functions of the mushroom bodies vary substantially between insect groups, particularly in the relative importance of visual and olfactory modalities (24). For example, in *Drosophila*, the mushroom bodies are important for olfactory learning (30, 31), but the mushroom body calyces (which are innervated by projection neurons carrying sensory information) receive comparatively less visual input (32) than other groups, including the apocritan Hymenoptera (33) and some Lepidoptera (27, 34), in which a large proportion of the calyx is innervated by projection neurons from visual centres. This variation suggests the mushroom body is evolutionarily labile in its response to differing ecological selective pressures, but few behavioural studies have investigated how behaviour varies with mushroom body size between closely related species.

Genetically determined increases in the size of key learning centres are unlikely to be the sole contributor to the emergence of behavioural differences. By definition, learning and memory require neural mechanisms that are environmentally sensitive. As such, neural plasticity and neural expansion may act as two, non-independent, axes underpinning cognitive evolution. Synaptic plasticity, which involves the reorganisation of synaptic connections between neurons, plays a key role in learning and memory (35, 36). The insect mushroom body exhibits a high degree of post-eclosion volumetric expansion (19, 37, 38), which often reflects changes in the number and density of microglomeruli (18, 39–43), synaptic complexes formed by connections between sensory projection neurons and Kenyon cells, the intrinsic neurons of the mushroom body (44). In contrast to vertebrates, where adult neurogenesis may play a role in some learning processes (45), many insects lack adult neurogenesis in the mushroom bodies, including honeybees (46), perhaps the most widely used insect model for studying the neural basis of advanced cognitive abilities (47). This suggests that synaptic plasticity may have pre-eminence over adult neurogenesis in the development of enhanced insect cognition. The combined effects of high interspecific variability in size and composition, and high intraspecific plasticity, make the mushroom bodies an excellent model for investigating the neural underpinnings of cognitive evolution. However, few studies have investigated interspecific variation in mushroom body plasticity and its potential role in cognitive evolution (48).

The Neotropical butterfly genus *Heliconius*, and their Heliconiini allies, are an emerging system for investigating the neuronal basis of interspecific cognitive variation. *Heliconius* have markedly expanded mushroom bodies relative to other Heliconiini (27–29) (Fig. 1), with this size increase primarily driven by increased neuron number and volumes of calyx receiving innervations from visual, rather than olfactory, processing centres (27). This expansion event occurred relatively recently (∼12-18 Ma) (49) and co-occurs with the dietary innovation of adult pollen feeding and associated derived foraging behaviours (50, 51) (Fig. 1). *Heliconius* establish “traplines”, long-term, stable foraging routes along which specific plants are regularly visited, suggesting an enhanced capacity for long term, visually-oriented spatial memory (52–54). Indeed, *Heliconius* have been shown experimentally to solve spatial learning assays across a range of spatial scales (Moura et al. *in press*), and individuals that are experimentally displaced by several hundred metres are able to return to their original locations (55), suggesting active site fidelity.

**Fig. 1.**
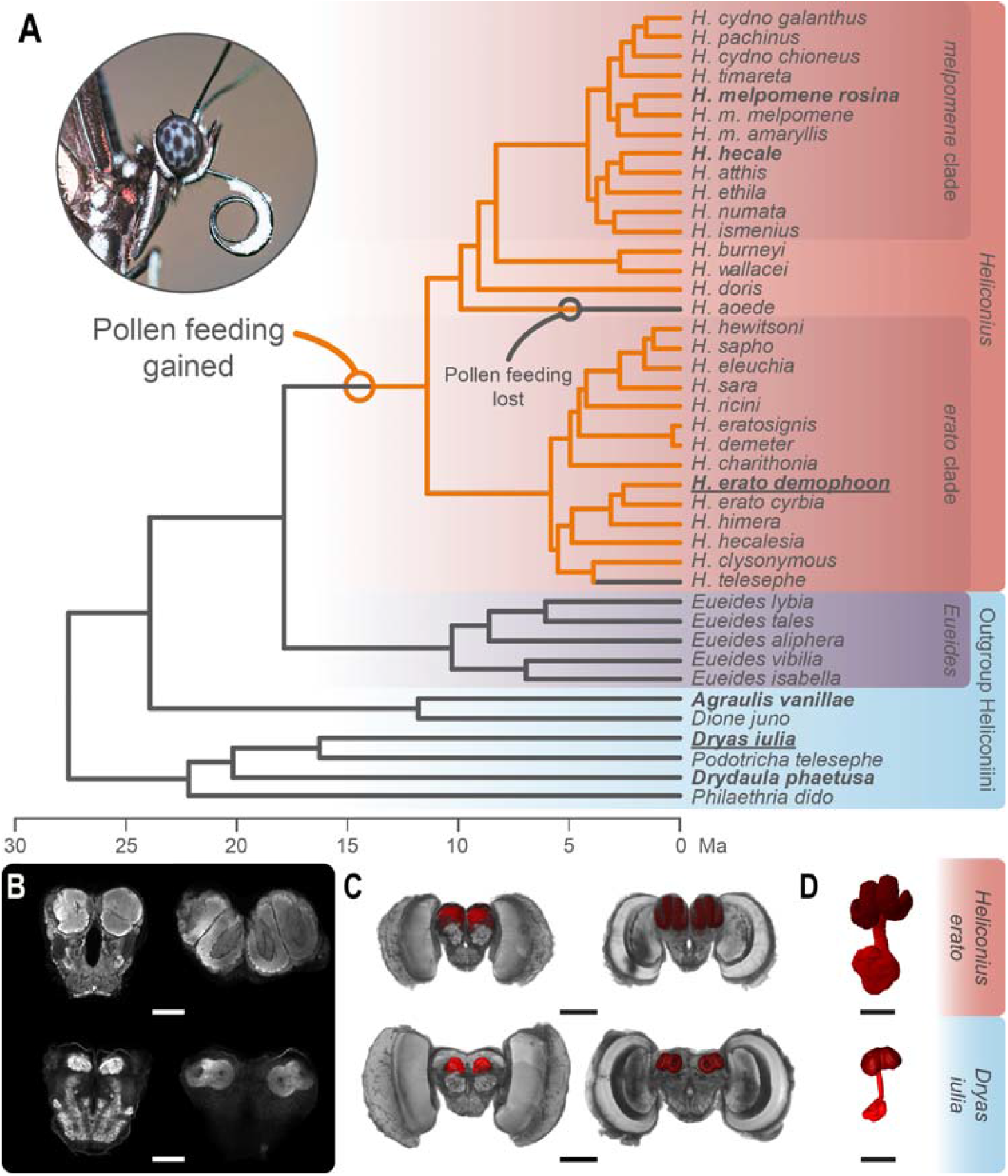
(*A*) Distribution of pollen feeding in Heliconiini butterflies with dated phylogeny adapted from (27). (*B*-*D*) Mushroom bodies in *Heliconius hecale* and *Dryas iulia*. (*B*) Confocal scans showing the anterior (left) and posterior (right) of the central brain (scale bars = 250 _μ_m). (*C*) 3D-reconstructions of the whole brain showing anterior (left) and posterior (right) with the mushroom bodies (red) (scale bars = 500 _μ_m). (*D*) Isolated 3D-reconstructions of the mushroom body showing the calyx (dark red) and peduncles and lobes (light red) scale bars = 250 μm).

Pollen feeding provides an adult source of amino-acids, and has co-evolved alongside a major increase in lifespan and reproductive longevity (56). This expanded adult period likely increases the benefit of long-term memories. Traplines can be followed faithfully for months (54), well past the maximum longevity of non-pollen feeding relatives (56). Consistent with this observation, comparative experiments have shown that *Heliconius erato* outperforms the Heliconiini *Dryas iulia* in their ability to learn and recall complex visual cues and maintain long-term memories of learned colour associations (27), indicative of substantial cognitive variation between *Heliconius* and other Heliconiini. Given their established role in long-term memory (39, 40, 57) and visual learning (18, 58–60) in other insects, changes in long-term memory may be intimately linked to mushroom body expansion in *Heliconius* butterflies.

Here, we extend previous data by testing visual long-term memory across six species, three *Heliconius* (*Heliconius melpomene* and *Heliconius hecale*) and three outgroup Heliconiini (*Agraulis vanillae* and *Dryadula phaetusa*), demonstrating a consistent, superior performance across the *Heliconius* genus. We subsequently investigate the neural basis of this behavioural difference, examining calyx volume, synapse density and the number of Kenyon cells in *Heliconius erato* and *Dryas iulia* that participated in the long-term memory assay, in conjunction with age-matched controls, and freshly eclosed butterflies. We quantify neural plasticity in these individuals, and test for neural correlates of learning experience.

## Results and discussion

### *Heliconius* show superior visual long-term memory relative to outgroup Heliconiini

We trained individuals from three *Heliconius* and three outgroup Heliconiini species to associate a food reward with either yellow or purple feeders over four days (Figs. S1, SS). *Heliconius* exhibited a slight increase in accuracy over non-*Heliconius* individuals (Fig. 2; z ratio=-2.240, P=0.048), but all six species successfully learned the association (Fig. 2, Table S4). We then tested the ability of butterflies to recall this learned association after a period of 8 days without these colour cues, during which they were fed solely on white feeders. *Heliconius* individuals exhibited markedly greater fidelity to the trained cues in a subsequent recall test (Fig. 2; z ratio=-5.807, P<0.0001). Furthermore, while all species exhibited a decline in accuracy over the eight days between the initial trained test and the long-term memory test (Fig. 2, Table S4), this drop-off was significantly higher for non-*Heliconius* individuals than in *Heliconius* (χ^2^=5.309, d.f.=1, P=0.0212). We further assayed the recall abilities of *H. melpomene, H. hecale, Agraulis vanillae* and *Dryadula phaetusa* after an additional four days deprived of the learned colour cues (a total of 13 days without reinforcement), again finding higher accuracy in the *Heliconius* individuals (Fig. 2; z ratio=-3.731, d.f.=inf, P<0.001). At this point, the colour preferences of *A. vanillae* and *D. phaetusa* were not different from random (Table S6), suggesting a loss of the learnt association. In contrast, *H. melpomene* maintained their learned preference during this period, and *H. hecale* exhibited a bias for the learned colour that approached significance (Table S6). To our knowledge, this period of 13 days represents the longest example of the persistence of a learned association without reinforcement in an insect, ahead of an 11-day period tested in the honeybee (61).

**Fig. 2.**
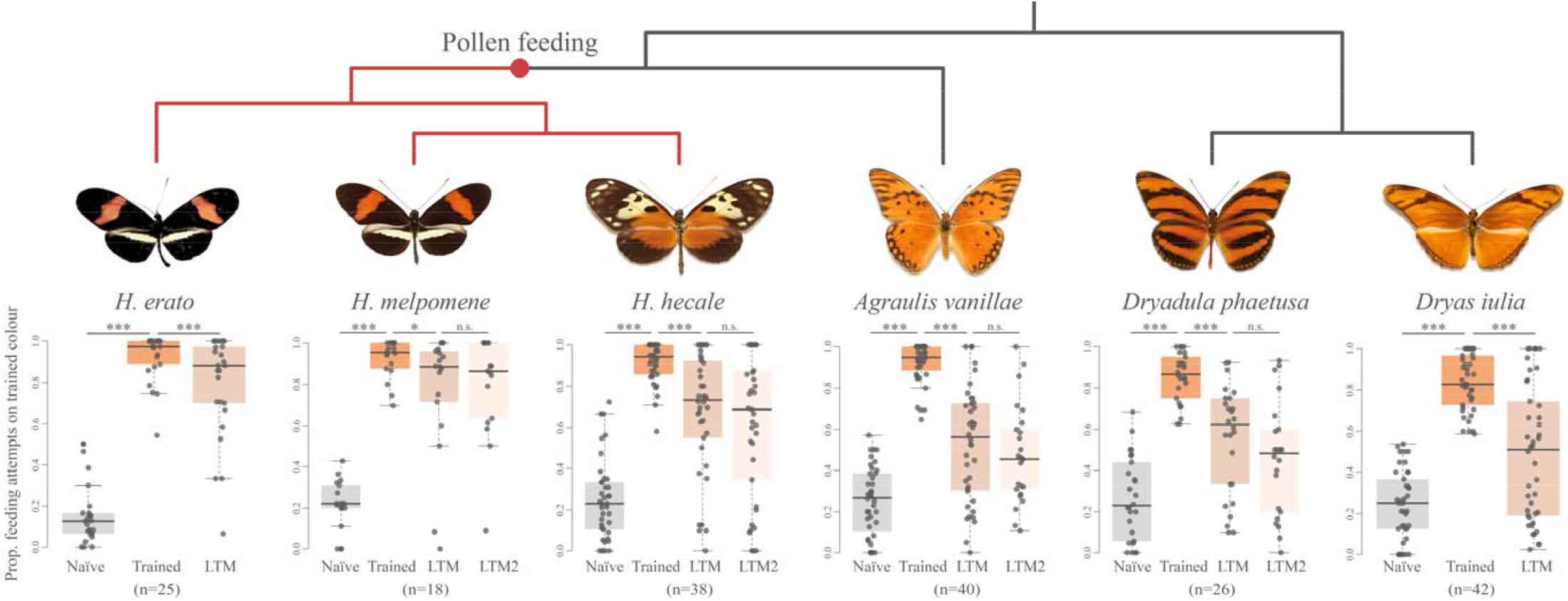
Long-term memory accuracy in six Heliconiini species in a two-colour preference assay. Trained = 16-hour recall performance after four days training. LTM = recall performance after eight days deprived of the learned colour stimuli by being fed solely on neutral (white) feeders; LTM2 = recall performance after an additional four days on white feeders. n.s. = no statistically significant difference; * = P<0.05; *** = P<0.001. Butterfly images from (83).

Our results reflect an important expansion previous data showing that *Heliconius erato* outperforms *Dryas iulia* in this task. By including two additional *Heliconius* species and two additional outgroup species, we sample major clades within *Heliconius* and across the Heliconiini, to confirm consistent shifts between *Heliconius* and other Heliconiini. These data strongly suggest that *Heliconius* as whole possess an enhanced visual long-term memory ability relative to other Heliconiini. This difference is consistent with mushroom body expansion in *Heliconius* facilitating an improved visual long-term memory, further supported by the dominant role of increased visual processing in *Heliconius* mushroom body expansion (27). This behavioural change may have been driven by the cognitive demands of traplining for pollen in the context of increased individual longevities. The role of the mushroom bodies in the formation and maintenance of olfactory long-term memories is well established (39, 57, 62). However, the present results add to evidence that for at least certain groups of insects, including many Hymenoptera (43), the mushroom bodies also play a significant role in visual long-term memory, mirroring the relative extent of visual and olfactory neural input (23, 24). In contrast to the significant clade effects we find in our long term memory assays, comparisons across a similar sample of Heliconiini species did not find *Heliconius* to be superior at learning shape cues (63) or reversal learning of colour cues (64). When coupled with our present results, these findings suggest that mushroom body expansion in *Heliconius* has not led to an overall improvement in general cognition, but rather enhancement in specific, ecologically-relevant cognitive tasks.

### Mushroom bodies of *Dryas iulia* and *Heliconius erato* vary in their response to learning experience

We further explored the neural underpinnings of this cognitive shift in *Heliconius* at the cellular and synaptic level by measuring several traits in the mushroom body calyx of *Heliconius erato* and *Dryas iulia* that had completed the long-term memory assay (Learning group). These individuals were compared with age-matched individuals from a “non-learning” environment (Control group) and freshly eclosed butterflies (Day 0 group). The “non-learning” group experienced the same experimental set up as the learning group, but both colours were equally reinforced and punished, preventing a conditioned preference (Figure S7, Table S7). Overall, when comparing Day 0 butterflies to either the Control or Learning groups, the mushroom bodies of *Heliconius erato* showed considerably more plasticity than *Dryas iulia* (Fig. 3, Tables S8 & S9). In *Heliconius erato*, the Learning and Control group individuals had a lower synapse density and fewer total synapses in the calyx, but increased calyx volumes, relative to Day 0 individuals (Fig. 3 *A*-*C*, Table S9). In contrast, for *Dryas iulia*, although similar trends were observed, neither synapse density, calyx volume, synapse number nor Kenyon cell number, varied significantly between groups (Fig. 3, Table S9). The age-associated increases in calyx volume, and decreases in synapse density, that we observe in *H. erato* have similarly been described in a number of Hymenoptera including bumblebees (18, 65), honeybees (41, 66), ants (42, 48) and paper wasps (67, 68). The age-dependent decrease in calyx synapse number observed in *Heliconius erato* is also consistent with observations in honeybees (41, 66, 69) and desert ants (42). This “pruning” of synapses, which is also present in vertebrates, is a recognised method of refining neural connectivity involving the selective elimination of axonal branches and increasing the strength of remaining synaptic connections (70).

**Fig. 3.**
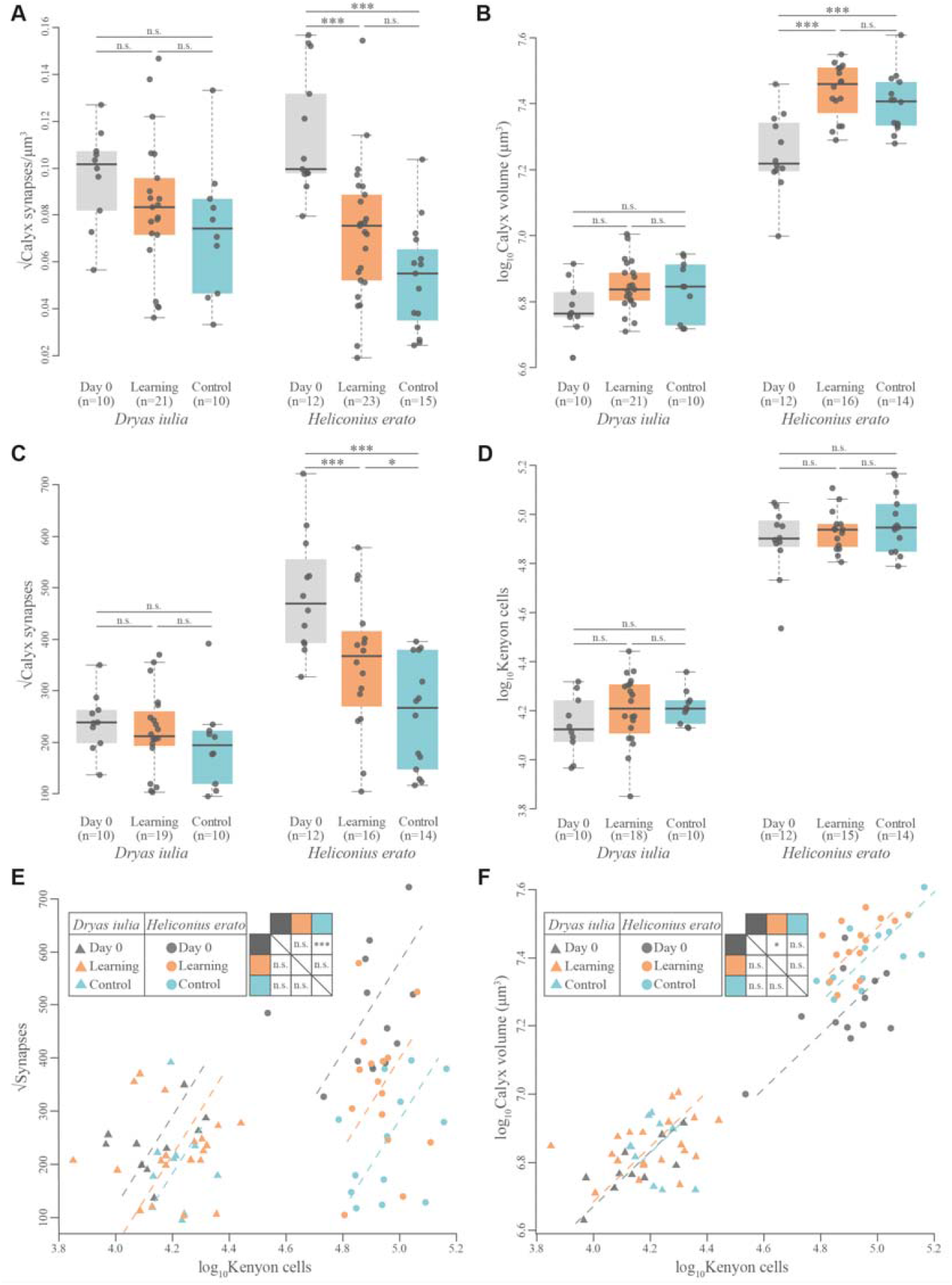
Variation in (*A*) synapse density in the calyx, (*B*) mushroom body calyx volume, (*C*) total synapse number in the calyx, and (*D*) total Kenyon cell number, between three treatment groups, Day 0, Learning and age-matched Controls, in *Dryas iulia* and *Heliconius erato*. (*E*),(*F*) Variations in elevation in the scaling relationship between Kenyon cell number and (*E*) number of calyx synapses, and (*F*) calyx volume across treatment groups treatment groups. Inset shows pairwise comparisons between groups with *Heliconius erato* above the diagonal and *Dryas iulia* below. n.s. = no statistical significant difference; * = P<0.05; ** = P<0.01; *** = P<0.001.

Notably, individual *Heliconius erato* in the Learning group had a significantly higher number of synapses in the calyx than the Control group individuals (Fig. 3 *C*, Table S9), which have similar synapse counts to *Dryas iulia* despite having much larger calyces (Fig. 3, Table S9). This suggests a higher maintenance of synapses in the Learning group in *Heliconius erato* and more extensive synaptic pruning in the Control group. The experience of the Learning and Control groups was identical, except for the reinforcement of colour cues during the four-day training period, a relatively modest environmental difference. Remarkably, these differences persisted for eight days after exposure to coloured feeders, suggesting that learning-induced synaptic connections in the calyx are being maintained for considerable amounts of time after exposure. Increased density or number of mushroom body synapses has previously been linked to visual (18) and olfactory (39, 40) learning and long-term memory in Hymenoptera, and these results extend those findings to visual learning and memory in a *Heliconius* butterfly. Interestingly, in *Drosophila*, synaptic reorganisation of the mushroom body appears to be essential for only certain associative learning tasks, being necessary for aversive, but not appetitive, associative odour learning (71).

*Heliconius erato* did not show an increase in calyx volume in response to associative colour learning specifically (Fig. 3 *B*), suggesting learning and memory of the colour cues was achieved solely through synaptic reorganisation. This mirrors findings in honeybees (39) and leaf-cutting ants (40), where olfactory learning was linked to increased synapse density without expansion of the calyx. Calyx growth has, however, been linked to visual learning in honeybees (18) and host plant learning in the butterfly *Pieris rapae* (19). In addition, consistent with previous findings (Alcalde et al., *in review*), we found no intraspecific group differences in Kenyon cell number, suggesting an absence of adult neurogenesis (Fig. 3, Table S9). Differences in synapse count between *Heliconius erato* groups therefore appear to be a result of changes in the number of synapses per Kenyon cell (Fig. 3 *E*, Table S10). Day 0 individuals had significantly more synapses per Kenyon cell than Control individuals, but not the Learning group (Fig. 3 *E*, Table S10). Learning group individuals also maintain significantly higher numbers of synapses per Kenyon cell compared to the Control group, when not correcting for multiple comparisons (Fig. 3 *E*, Table S10). Together with the lack of neurogenesis in honeybees (46), the present findings suggest that, in insects, sophisticated cognitive ability can be achieved without adult neurogenesis, despite its importance in some vertebrate brain regions (45).

### Recall accuracy in *Heliconius erato*, but not *Dryas iulia*, is associated with increased synapse density and number in the mushroom body calyx

We further tested for specific neural correlates of within-species variation in recall performance in both the initial recall (16 hours removed from the colour cues) and long-term recall tests (eight days removed). For *Heliconius erato*, performance in the initial recall test was positively correlated with calyx synapse density and number, and the ratio of synapses to Kenyon cells (Fig. 4 *G I K*, Table S10). Together with our prior finding of learning experience being associated with an increased calyx synapse count in *Heliconius erato*, this strongly suggests that the synaptic organisation of the calyx plays a role in visual learning and memory in *Heliconius*. These results also mirror data from honeybees where a similarly positive correlation is reported between synapse density in the mushroom body collar, an area of the calyx receiving solely visual input, and recall accuracy (26). Unlike *Heliconius erato*, initial recall accuracy performance in *Dryas iulia* was not correlated with calyx synapse density or number (Fig. 4 *A C E*, Table S10), but was, surprisingly, *negatively* correlated with calyx volume (Fig. 4 *B*, Table S10). This is suggestive of an altered relationship between mushroom plasticity and the consolidation of the learned food-colour association between these two species. Neither species showed any correlation between performance in the long-term recall test and measured traits in the calyx. However, when a clear outlier individual is removed, positive relationships with synapse count (χ^2^=9.263, d.f.=1, P=0.002) and the ratio of synapses to Kenyon cells (χ^2^=10.918, d.f.=1, P=0.001) were recovered in *H. erato* (Fig. S9), which may suggest the associations detected above persist to the 8-day recall trial. Regardless, the above results suggest an important role for synaptic reorganisation in the mushroom body calyx in the formation of visual associative memories in *Heliconius*.

**Fig. 4.**
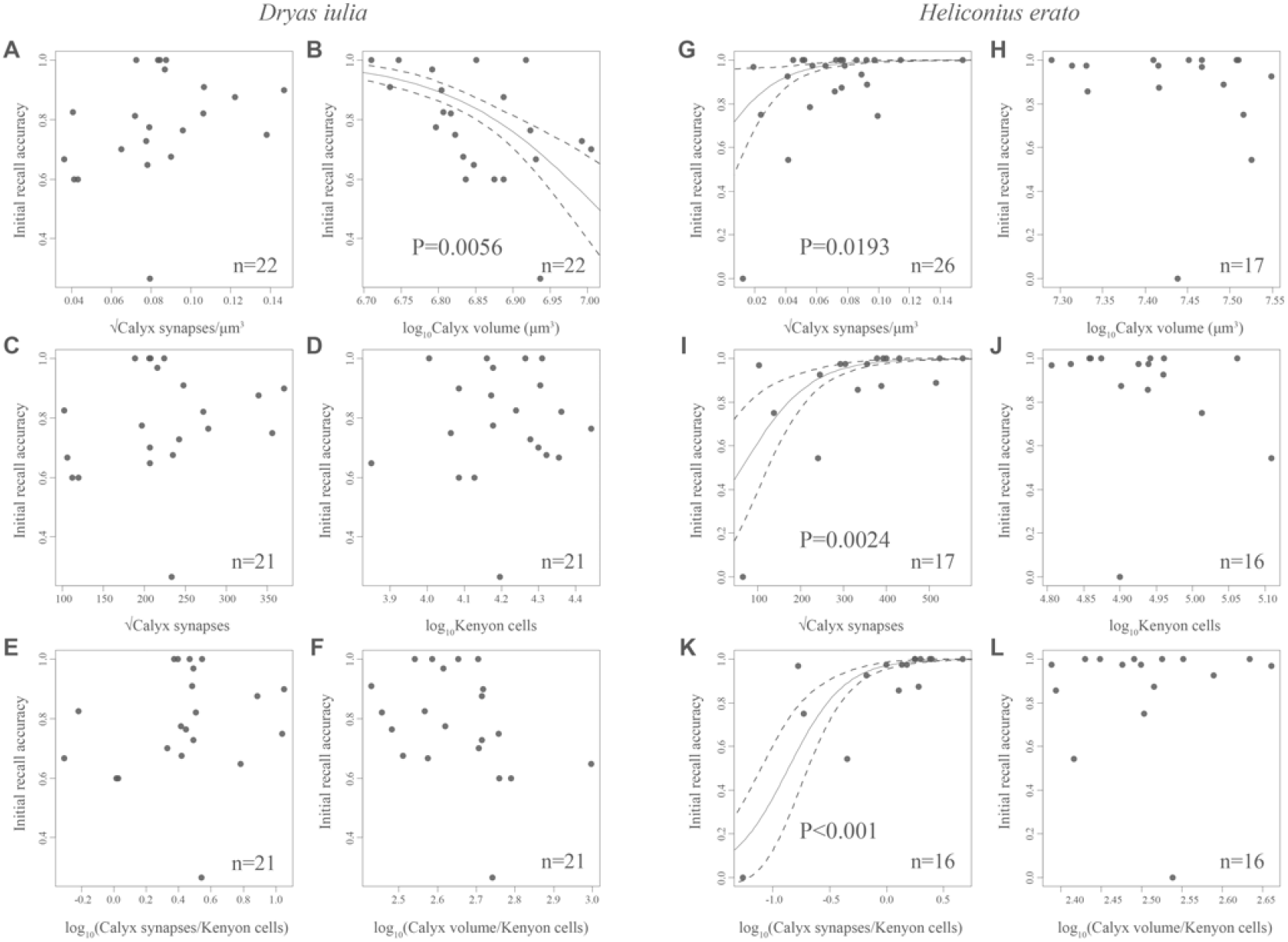
Relationship between 16-hour recall accuracy of a learned food-colour association in a two-colour preference test and several neuroanatomical measurements in the mushroom body calyx for (*A*)-(*F*) *Dryas iulia* and (*G*)-(*L*) *Heliconius erato*. Regression lines, with standard errors, are shown only when the trait was a significant predictor of performance.

## Summary

Our data strongly suggest that mushroom body expansion in *Heliconius* is associated with an improved long-term memory and positions *Heliconius* as an important example of structural changes in the brain supporting a cognitive shift over a relatively small phylogenetic scale (∼12-18 Ma) (49). These neuroanatomical and cognitive shifts coincide with the emergence of trapline foraging, which seemingly depends on, in part, the long-term memory of resource locations, providing a likely selective pressure driving these changes (50). We provide evidence that the mushroom bodies of *Heliconius erato* are not only larger than those of the closely-related Heliconiini *Dryas iulia*, but exhibit greater synaptic restructuring in response to experience. In *Heliconius erato*, visual learning experience, was associated with an increased retention of synapses in the calyx, while recall accuracy of the learned colour-cue association after 16 hours was positively correlated with calyx synapse density and number. These results point to synaptic connections between Kenyon cells and projection neurons from primary visual neuropils playing a key role in visual learning and memory in *Heliconius*, mirroring findings in honeybees (18). In a further convergence with honeybees (46), we found no evidence of increased Kenyon cell number, suggesting learning and memory in *Heliconius* is achieved primarily through synaptic reorganisation. Mushroom body expansion in *Heliconius* and Hymenoptera therefore appears convergent in multiple developmental and anatomical traits.

In summary, we have combined behavioural data with large- and fine-scale neuroanatomical measurements in an interspecific, comparative framework, to investigate the neural underpinnings of interspecific variation in cognition. This is, to our knowledge, the first study to present measurements of both cognitive performance and cellular organisation of the mushroom body across two closely-related insect species. In doing so, we show that volumetric expansion of a specific brain region, and increased synaptic plasticity, coincided with a discrete, behavioural shift, providing an important case study towards understanding the evolution of cognition.

## Methods

### Animals

All experiments used freshly eclosed butterflies reared from stock populations established with locally-caught, wild butterflies at the Smithsonian Tropical Research Institute in Gamboa, Panama. The long-term memory assay used three species of *Heliconius* (*H. erato, H. hecale* and *H. melpomene*) and three species of non-*Heliconius* Heliconiini (*Dryas iulia, Dryadula phaetusa* and *Agraulis vanilla*). *H. erato* and *Dryas iulia* individuals from this experiment were further dissected for the neuroanatomical investigations.

### Long-term memory assay

We tested the ability of Heliconiini butterflies to remember a learned food-colour association after eight days without reinforcement. Freshly eclosed individuals were first subject to two days of pre-training, being fed solely on white feeders containing a sugar-protein solution (20% sugar, 5% Vertark Critical Care Formula, 75% water, w/v). Butterflies were then tested for their initial colour preference by counting their feeding attempts on empty purple and yellow feeders over a four hour period. Butterflies were then trained to associate a food-reward with the non-preferred colour for four days. Rewarded feeders were filled with a sugar-protein solution and non-rewarded feeders contained a saturated quinine solution as an aversive stimulus. Colour preferences were then re-tested to verify individuals had learned then colour-food association. Butterflies were then fed solely on neutral (white) feeders for eight days, before their colour preference was retested. *Heliconius erato* and *Dryas iulia* individuals were then sacrificed for dissection, while *Heliconius melpomene, Heliconius hecale, Agraulis vanillae* and *Dryadula phaetusa* were fed for a further four days on white feeders and then subject to a final colour preference test.

### Immunohistochemistry and brain imaging

We compared the mushroom body volume and structure of *Heliconius erato* and *Dryas iulia* individuals from three groups: (1) individuals that participated in the long-term memory assay (Learning group); (2) individuals age-matched to the Control group and exposed to a “non-learning” environment, where purple and yellow feeders were equally rewarding as punishing, for four days, before being fed on white feeders for eight days (Control group), and; (3) individuals dissected the day of eclosion (Day 0).

Prior to dissection brains were fixed in zinc-formaldehyde solution. We used mono-clonal anti-synapsin antibody (mouse anti-SYNORF1: 3C11, DSHB, RRID:AB_2315424) and Cy2-conjugated secondary antibody (Cy2 goat anti-mouse IgG: 115-225-146, Jackson ImmunoResearch, RRID: AB_2307343) anti-mouse antibody to mark synapses in the calyx. A DAPI (D9542, Sigma-Aldrich) stain was used to identify Kenyon cell bodies nuclei (72, 73). HRP (Rabbit anti-horseradish peroxidase, P-7899, Sigma-Aldrich, RRID:AB_261181) and Cy3-conjugating secondary antibody (Cy3 goat anti-rabbit IgG: 111-165-144, Jackson ImmunoResearch, RRID: AB_2338006) was used to label neural membranes to confirm that counted nuclei were neuronal (74). Prior to imaging, brains were sectioned into 80_μ_m slices to permit imaging at high magnification. All imaging was performed on a laser-scanning confocal microscope (Upright Leica SP5, Leica Microsystem) with a mechanical *z*-step of 1 _μ_m at a resolution of 1024×1024. The whole calyx was imaged using a dry 10X objective with a numerical aperture of 0.4 and confocal scans were segmented using Amira 3D 2021.2 (Thermo Fisher Scientific) to estimate its volume, and the volume of the surrounding Kenyon cell cluster. Kenyon cells and synapses were imaged using a 63X objective with a numerical aperture of 1.3, under glycerol immersion. For each individual, five 50×50×15_μ_m boxes were randomly placed in the calyx, within which synapse densities were estimated using the *3D Objects Counter* function in ImageJ (National Institutes of Health) (Fig. S4 *A, B*) (Alcalde et al., *in review*). Note, unlike Hymenoptera, the visual and olfactory regions of the Heliconiini calyx lack clear morphological boundaries, meaning we were unable to reliably place boxes specifically in visual or olfactory calyx. However, boxes were positioned in distal positions where possible to largely sample visual areas. Kenyon cell density was estimated from five randomly selected 25×25×15_μ_m boxes in the cell cluster, with cell numbers counted using the *Modular Image Analysis* (MIA) and *Stardist* plugins in ImageJ (75, 76) (Alcalde et al., *in review*).

## Statistical analyses

Performance in the long-term memory trials was analysed with generalised linear mixed models (GLMMs) using the *glmer* function from the *lme4* package v1.1-21 in R v 4.1.0 (77). All models used a binomial distribution with purple and yellow feeding choices as dependent variables and individual- and observation-level random effects included to account for overdispersion. Diagnostics for all models were assessed using the package *DHARMa* v0.4.4 (78). Post-hoc comparisons among relevant pairs of species, clades, or trials were made by obtaining the estimated marginal means using the package *emmeans* v1.7.0 (79) and were corrected for multiple comparisons using Šidák correction. Differences between *Heliconius* and outgroup Heliconiini were tested by including membership in the *Heliconius* genus as fixed effect with species as a random effect.

For the neuroanatomical comparisons, *Heliconius erato* and *Dryas iulia* synapse densities and counts were first square-root transformed, and calyx volume and Kenyon cell counts were log10 transformed to better fit a normal distribution. We then ran a series of generalised linear models (GLMs) with a Gaussian distribution using the *glm* function in R v 4.1.0 GLMs testing for differences in these neuroanatomical traits between groups within species and between species within groups. Variation in the scaling relationships between Kenyon cell number and number of synapses in the calyx, and Kenyon cell number and calyx volume, were tested using the *sma* function in the R package *smatr* v 3.4-8 (80). The “robust” option was set to true for these analyses and multiple comparisons were corrected for (81). Finally, we tested whether specific neural traits correlate with recall performance by running a series of binomial GLMMs using the function *glmmTMB* from the package *glmmTMB* v 1.1.2.3 for R (82), with the neural trait of interest as a fixed effect and ID as a random effect. These analyses were repeated for both the initial recall test and the long-term recall test.

## Acknowledgements

We thank the Ministerio del Ambiente, Panama, and the science support, administrative, and facilities staff at the Smithsonian Tropical Research Institute in Panama for making this research possible. In particular, we thank Oscar Paneso and Cruz Batista for help in butterfly stock and host plant care. We also gratefully acknowledge the Wolfson Bioimaging Facility for their support and assistance in this work. This work was supported by a NERC Independent Research Fellowship (NE/N014936/1) and an ERC Starter Grant (758508) to SHM, and a PhD Studentship from Trinity College, Cambridge to FJY.

