## Supplementary materials for "Enhanced long-term memory and increased mushroom body plasticity in *Heliconius* butterflies"

### Detailed methods

#### Animals

All experiments used freshly eclosed, captive-reared butterflies with no prior experience. Butterfly stock populations were established with locally-caught, wild butterflies and maintained at the insectaries at the Smithsonian Tropical Research Institute in Gamboa, Panama. Stock butterflies were kept in 2x2x3m mesh cages in ambient conditions with natural light. Larvae were reared in mesh pop-ups and were provided with fresh leaves daily. The long-term memory trials used three species of *Heliconius* (*H. erato*, *H. hecale* and *H. melpomene*) and three species of non-*Heliconius* Heliconiini (*Dryas iulia*, *Dryadula phaetusa* and *Agraulis vanilla*). *H. erato* and *Dryas iulia* individuals were dissected for the neuroanatomical investigations. *H. erato*, *Dryas iulia*, *Dryadula phaetusa* and *Agraulis vanillae* were reared on *P. biflora*, *H. melpomene* on *P. triloba*, and *H. hecale* on *P. vitifolia*. Training and testing of butterflies was conducted in 2x2x3m mesh cages in ambient conditions under natural light. A single *Psychotria elata*, with all flowers removed, was placed in the rear right corner of these cages as a roosting site.

#### Long-term memory assay

Long-term memory (LTM) experiments, using colour cues, were carried out on captive-reared butterflies between January and April 2019 in Gamboa, Panama. The experiments used two colours, purple and yellow, colours chosen based on previous experiments using *H. erato* which showed that neither colour was particularly attractive (1). The experiments used five-pointed, star-shaped, artificial feeders made from coloured foam, 3 cm in diameter, with a centrally placed 0.5 ml Eppendorf tube that could be filled with liquid (Fig. S1).

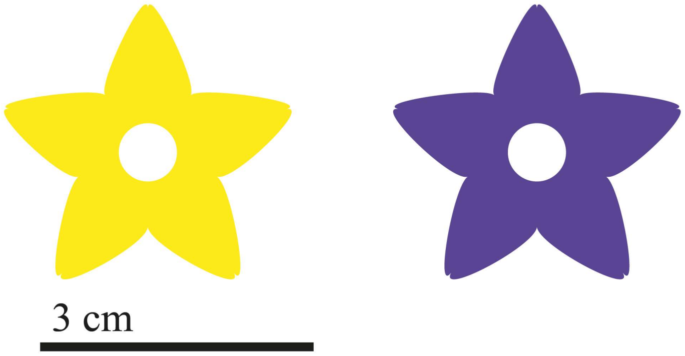


**Fig. S1** Purple and yellow feeders used for the long-term memory and reversal learning assays.

The day after eclosion, individuals were transferred to a pre-training cage, where they were fed solely with white artificial feeders containing a sugar-protein solution (20% sugar, 5% Vertark Critical Care Formula, 75% water, w/v) for two days (from 08:00 to 12:00) to familiarise them with the use of artificial feeders. After pre-training, butterflies were introduced to a testing cage to determine initial feeding preferences between purple and yellow. Testing cages contained 12 purple and 12 yellow feeders (Fig. S1) arranged randomly in a 4 X 6 grid, with 6.5 cm between feeders on each side. To ensure that butterflies responded exclusively to visual cues, feeders in the testing cages were empty. Preference testing lasted for four hours from 08:00 to 12:00 and was filmed from above using a GoPro Hero 5 camera mounted on a tripod. Butterflies were individually numbered on their wings for identification using a permanent marker. The film was then reviewed to count the number of feeding attempts per individual on each colour, with up to 40 attempts recorded per individual. A feeding attempt was only counted if the butterfly landed on the feeder and probed it with its proboscis.

Butterflies were then trained to associate a food reward with their non-favoured colour, based on the results of their initial preference test. For butterflies that initially preferred purple, the training cage contained yellow feeders containing a sugar-protein solution, and purple feeders containing a saturated quinine solution, an aversive stimulus. The opposite arrangement was employed for individuals that initially preferred yellow. This training period lasted for four full days (Fig. S2). After training, butterfly preferences were re-tested, following the same protocol as the initial preference test, to verify that individuals had indeed acquired the colour-food association. After the trained preference test, butterflies were placed for eight days in a cage identical to the pre-training cage, containing only white feeders filled with a sugar-protein solution. The deprivation of colour stimuli for this period allowed for testing the long-term memory retention of the colour-food association acquired during the training period, and ensured that long-term memory was being tested rather than short-term or mid-term memory (2). A period of eight days was chosen because *Heliconius* are known to maintain their foraging routes over periods ranging from weeks to months (3–6), during which time a pollen resource could be unproductive for several days due to competition or damage, but ultimately rewarding over the long term. Butterflies were then subject to a third preference test to determine if the learned preference was maintained, following the same protocol as the initial preference test (Figure S2). *H. melpomene*, *H. hecale*, *Dryadula phaetusa* and *Agraulis vanilla* individuals were also subject to an additional extended long-term memory test by placing them in a cage with only white for a further four days before a final preference test.

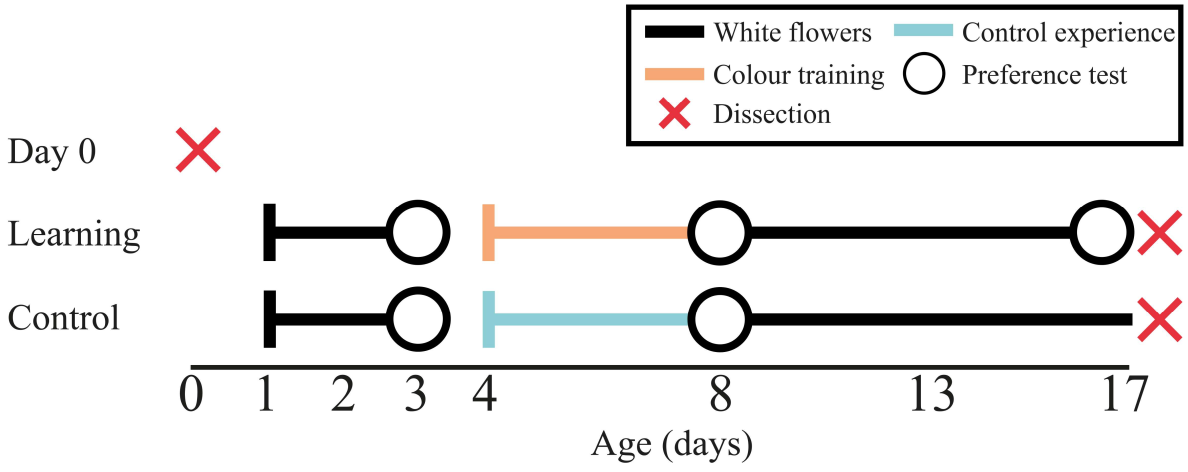


**Fig. S2** Schematic of the three time line for the long-term memory experiment, including the Control and Day-0 groups used for the neuroanatomical parts of the study. Only *Dryas iulia* and *Heliconius erato* individuals were dissected.

**Kenyon cell and synapse counting and calyx measurement**

*(a) Treatment groups*

Neuroanatomical measurements were taken for *Dryas iulia* and *Heliconius* erato from three treatment groups – Day 0, Learning and Control (Fig. S2). Day 0 butterflies comprised individuals that were dissected the evening of their day of emergence. These butterflies were kept in a small, mesh pop-up cage until dissection and had no foraging or free-flight experience. The Learning Group comprised *Dryas iulia* and *Heliconius erato* individuals that participated in the long-term memory experiment and were dissected the day they finished (Learning Group; Fig. S2). The Control Group consisted of individuals aged-matched to the Learning group reared in a “non-learning” environment. Control individuals were acclimatised to the use of artificial feeders and then tested for naïve colour preference following the same protocol used for the Learning Group. These butterflies were then introduced to a “training” cage for four days, which contained an even number of purple and yellow feeders, presented in random spatial arrangement. Unlike the Learning group, where feeder colour was consistently associated with either a food reward or quinine punishment, half of the feeders for each colour were filled with the rewarding sugar-protein mixture, and half with the aversive quinine solution. Each colour, therefore, was as equally rewarding as punishing. After four days in this environment, colour preference was tested again following the protocol established for the long-term memory experiment. The Control butterflies were then introduced to a cage with white feeders for eight days, replicating the long-term memory waiting period of the Learning group; Fig. S2). Finally, after eight days feeding on white feeders, Control butterflies were exposed to empty purple and yellow feeders from 08:00 to 12:00 on their final morning, mirroring the last preference test of the Learning Group, before being dissected in the evening.

*(b) Brain dissection, fixing and training*

All brains were dissected and fixed at the Smithsonian Tropical Research Institute in Gamboa, Panama, following established protocols (7, 8). Butterflies were decapitated using scissors and the head was submerged under HEPES-buffered saline (HBS; 150 mM NaCl; 5 mM KCL; 5 MM $CaCL_2$; 25 mM sucrose; 10 mM HEPES; pH 7.4). A small aperture was cut into the head cuticle between the eyes to improve permeation of the fixative. The brain was fixed *in situ* for 16-20 hours at room temperature under gentle agitation in zinc-formaldehyde solution (ZnFA; 0.25% [18.4 mM] $ZnCl_2$; 0.788% [135 mM] NaCl; 1.2% [35 mM] sucrose; 1% formaldehyde). After fixing, the whole brain was dissected out under HBS using a scalpel and forceps, placed into 80% methanol, 20% dimethyl sulfoxide (DMSO) under agitation for 2 hours and then transferred to 100% methanol for long-term storage at -20°C.

Brains were subject to three types of staining (9). Anti-synapsin was used to mark synapses in the calyx (10–12). We used DAPI to identify nuclei in Kenyon cell bodies (8, 13), along with HRP (anti-horseradish peroxidase) to label neuron membranes to confirm that counted nuclei were neuronal (14). Brains were stained in batches of eight that included individuals from all treatment groups to avoid the possibility of batch effects skewing results. Prior to staining, brains were first rehydrated in a decreasing methanol series (90%, 70%, 50%, 30%, 0% in 0.1 M Tris buffer, pH 7.4) for 10 minutes each. Because quantification of the Kenyon cells and synapses requires imaging the brain at 63X magnification, it was necessary to section the brains so that the calyx tissue would be within the working distance of the objective lens of the microscope. Brains were embedded in 5% agarose which was cut into a rectangular prism. The agarose block was cut along a corner so that individual slices could be correctly orientated later during mounting. The embedded brain was submerged in 0.1 M Tris buffer and sliced horizontally into 80 μm sections using a Leica VT1000 S vibrating blade microtome.

After sectioning, brain slices were blocked in PBSd-NGS (1% DMSO, 5% normal goat serum (NGS) diluted in 0.1 M phosphate-buffered saline (PBS; 7.4 pH)) for two hours. A mono-clonal antibody targeting synapsin (mouse anti-SYNORF1: 3C11, DSHB,

RRID:AB_2315424; [1:30]) and HRP (Rabbit anti-horseradish peroxidase, P-7899, Sigma-Aldrich, RRID:AB_261181; [1:5000]) were diluted in PBSd-NGS, then applied at a 1:30 dilution in PBSd-NGS and kept for three days at 4°C under low agitation. Samples were then washed in PBS (3 x 2 hours) before applying the Cy2-conjugated secondary antibody (Cy2 goat anti-mouse IgG: 115-225-146, Jackson ImmunoResearch, RRID: AB_2307343; [1:100]) and Cy3-conjugating secondary antibody (Cy3 goat anti-rabbit IgG: 111-165-144, Jackson ImmunoResearch, RRID: AB_2338006; [1:200]), in PBSd-NGS. Samples were then kept at 4°C under low agitation for a further three days. Samples were then rinsed in PBS (3 x 2 hours). In preparation for the DAPI stain, samples were washed in 0.2% Triton in distilled $H_2O$ for 10 minutes. DAPI was applied 1:1000 in 0.2% Triton and $H_2O$ under agitation for 30 minutes at room temperature. After, samples were permeabilised in 0.2% Triton and $H_2O$ for 10 minutes and then in 0.2% Triton and PBS (4 x 10 minutes) and then placed overnight in 60% glycerol in PBS. Brain sections were then mounted in 80% glycerol on slides under a coverslip sealed with nail polish and stored in the dark. Due to the delicate nature of the 80 μm slices, samples from some individuals suffered physical damage during staining and mounting meaning that their total calyx volume, and synapse and Kenyon cell counts could not be reconstructed.

#### *(c) Confocal microscopy and image processing*

All brains were imaged using a laser-scanning confocal microscope (Upright Leica SP5, Leica Microsystem, Mannheim, Germany), at a resolution of 1024x1024 pixels. Mushroom body calyces were scanned using 10X dry objective (0.4 NA), with a mechanical *z*-step of 1 μm, with each brain section scanned individually. Kenyon cells and synapses were imaged using the 63X objective (1.3 NA) under a glycerol immersion, with a mechanical *z*-step of 1 μm. For each individual, five regions of mushroom body calyx and Kenyon cell cluster were chosen at random for scanning. Cy2 was excited with the Argon laser at 488 nm. The solid-state lasers were used to excite DAPI at 405 nm and Cy3 at 561 nm. Wavelengths were scanned sequentially and received on photomultiplier tubes.

Brain images were processed using ImageJ v 1.53n and Amira 3D 2021.2. Calyx volumes were reconstructed using Amira, with each brain section segmented separately. For each stack, every third or fourth image was manually segmented by highlighting the region covered by the calyx and then interpolated across the *z*-dimension (Fig. S3). The *measure statistics* function was used to extract volumes (in $\mu m^3$) for each section, which were later added together for the total calyx volume and a correction factor of 1.85 was applied. Synapse densities were estimated using ImageJ. Following protocols established in (9), the *3D Objects*

*Counter* function was used to automatically count 3D objects within five 50x50x15µm boxes (Fig. S4 *A*, *B*). For each scan, the brightness threshold was adjusted manually so that only distinct objects were counted. To reduce noise, objects smaller than 10 voxels were not counted. Note, unlike Hymenoptera, the visual and olfactory regions of the Heliconiini calyx lack clear morphological boundaries, meaning we were unable to reliably place boxes specifically in visual or olfactory calyx. However, boxes were positioned in distal positions where possible to largely sample visual areas. The total number of synapses in the calyx was then estimated by calculating the average synapse density across the five scans and multiplying it by the total calyx volume, assuming homogeneity of synapse densities across the calyx. Kenyon cell cluster volumes and total Kenyon cell numbers were determined in a similar manner. Kenyon cell density was estimated from five randomly selected 25x25x15µm boxes in the cell cluster (Fig. S4 *C*, *D*). Cell numbers within each box were automatically counted using the *Modular Image Analysis* (MIA) and *Stardist* plugins in ImageJ (15, 16)(Alcalde et al., *in review*). *Stardist* detects objects with star-convex shape priors and can be used for detecting cells. Total cell counts were then estimated by multiplying the average density by the total volume of the cell cluster.

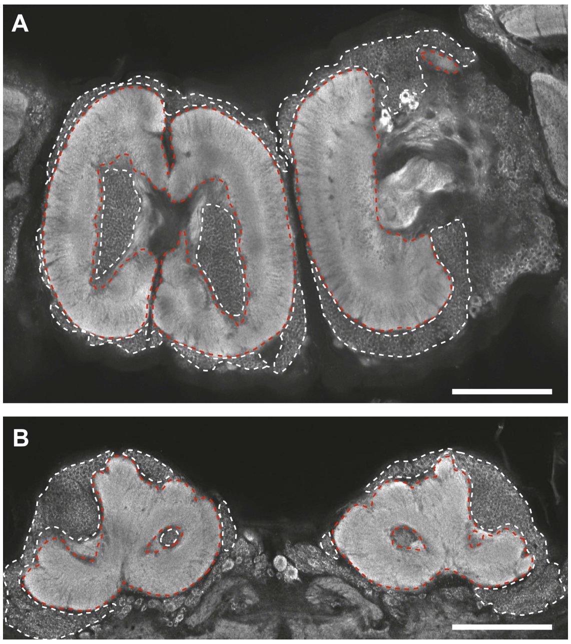


**Fig. S3** Examples of HRP staining of the mushroom body calyx, imaged at 10X using a laser-scanning confocal microscope, in (*A*) *Heliconius erato* and (*B*) *Dryas iulia* from the Learning group. White dashed line shows the calyx. Red dashed line shows the Kenyon cell cluster. Scale bar represents 200 µm.

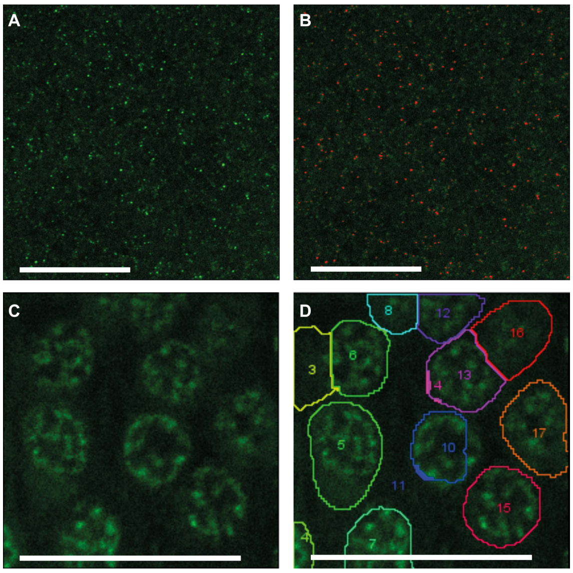


**Fig. S4** (*A*),(*B*) Synapses and (*C*),(*D*) Kenyon cells in the mushroom body calyx of *Heliconius erato*, imaged at 63X with a laser-scanning confocal microscope. (*A*) Synapses revealed through anti-synapsin staining, which are (*B*) automatically counted using *3D Object Counter* in ImageJ. (*C*) Kenyon cell nuclei revealed through DAPI staining which are (*D*) automatically counted using *Modular Image Analysis* and *Stardist* in ImageJ. Scale bar represents 20 µm.

### Statistical analyses

Performance in the long-term memory trials was analysed with generalised linear mixed models (GLMM) using the *glmer* function from the *lme4* package v1.1-21 in R v 4.1.0 (17). All models used a binomial distribution with purple and yellow feeding choices as dependent variables and individual-ID was included as a random effect. Post-hoc comparisons among relevant pairs of species, clades, or trials were made by obtaining the estimated marginal means using the package *emmeans* v1.7.0 (18) and were corrected for multiple comparisons using Šidák correction. To test for interspecific differences in performance, species was included as a fixed effect. Differences between *Heliconius* and outgroup Heliconiini were tested by including membership in the *Heliconius* genus and trial as fixed effects with species as a random effect. To test for potential differences between *Heliconius* and non-*Heliconius* in the drop in performance between the initial preference test and the long-term memory test, an interaction between *Heliconius*-membership and trial was included. To account for overdispersion, an observation-level random effect was included. Significant deviation from random colour preference during the initial preference test and the second long-term memory test was assessed using a null generalised linear mixed model. Diagnostics for all models were assessed using the package *DHARMa* v0.4.4 (19). For both the long-term memory and reversal learning trials, individuals that exhibited less than 50% accuracy during the initial trained preference test, and therefore did not appear to have learned the food-colour association, were removed from the dataset. In total, 9 individuals were so removed from the dataset: 1 out of 26 *H. erato* individuals were removed, in addition to 2 of 20 *H. melpomene*, 2 of 40 *H. hecale*, 2 out of 42 *Agraulis vanillae*, 1 of 42 *Dryas iulia*, 1 of 27 *Dryadula phaetusa*. A generalised linear model treating species as a fixed effect did not show significant variation between species in the proportion of individuals scoring less than 50% during the first training test ($\chi^2$=1.514, d.f.=5, P=0.911).

*Heliconius erato* and *Dryas iulia* synapse densities and counts were first square-root transformed, and calyx volume and Kenyon cell counts were $\log_{10}$ transformed to better fit a normal distribution. We then ran a series of generalised linear models (GLMs) with a Gaussian distribution using the *glm* function in R v 4.1.0 GLMs testing for differences in these neuroanatomical traits between groups within species and between species within groups. Species and group, and their interaction, were, thus, included as effects. For these analyses, we removed two individuals from the Learning group, one from each species, whose accuracy was less than 50% during the initial recall test, on the basis that those individuals had not acquired a learned association with the trained colour. All post-hoc comparisons were made by obtaining the estimated marginal means using the package *emmeans* v1.7.0 (18) and were corrected for multiple comparisons using the Šidák correction. We further tested for variation in the scaling relationships between Kenyon cell number and number of synapses in the calyx, and Kenyon cell number and calyx volume, using the *sma* function in the R package *smatr* v 3.4-8 (20). This analysis allows for the detection of shifts in elevation in the scaling relationship between two traits. The “robust” option was set to true for these analyses and multiple comparisons were corrected for (21). Finally, we tested whether specific neural traits correlate with recall performance in the initial recall test and long-term memory test. For each species, we ran a series of binomial GLMMs using the function *glmmTMB* from the package *glmmTMB* v 1.1.2.3 for R (22), with the trait of interest as a fixed effect and ID as a random effect. These analyses were repeated for both the initial recall test and the long-term recall test.

### Supplementary results

#### Interspecific variation in initial colour preference

Species varied significantly in their naïve colour preferences (Fig. S5, $\chi^2$=33.968, d.f.=5, P<0.0001). There were no significant differences between the naïve preferences of *H. melpomene*, *H. hecale*, *Dryadula phaetusa* and *Dryas iulia* (Fig. S5, Table S1), which did not significantly differ from 50% (Table S2). However, the naïve preferences of *Agraulis vanillae* and *H. erato* significantly differed from the other species and were biased towards purple over yellow (Fig. S5, Table S1, Table S2).

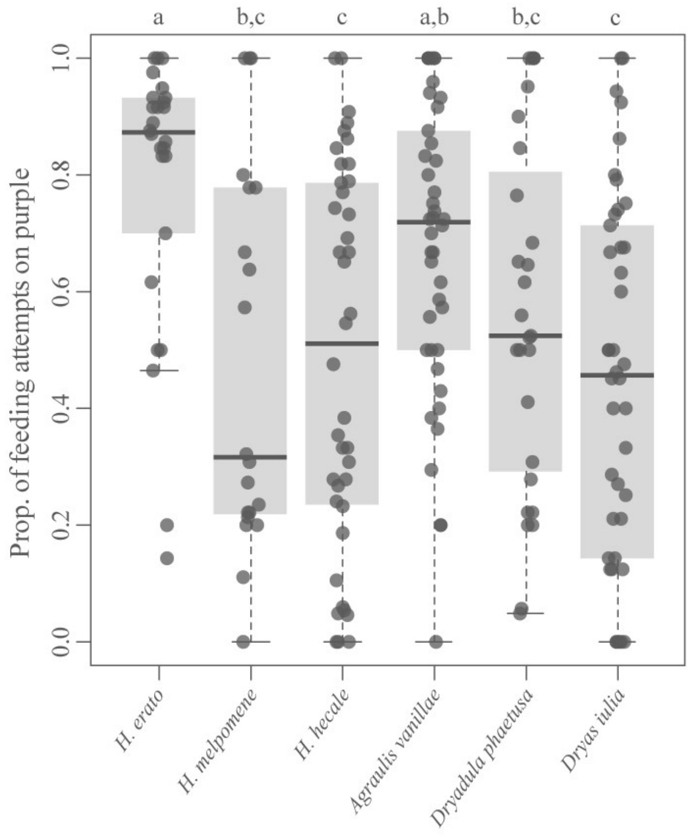


**Fig. S5** Naïve colour preference between purple and yellow for three *Heliconius* species (*H. erato* (n=26), *H. melpomene* (n=20) and *H. hecale* (n=40)) and three non-*Heliconius* Heliconiini, *Agraulis vanillae* (n=42), *Dryadula phaetusa* (n=27) and *Dryas iulia* (n=42) used in the long-term memory experiment. Notation at the top shows significantly different pairwise comparisons based on Table S1.

**Table S1** Pairwise comparisons between species in naïve preference between purple and yellow feeders. Bottom left shows z-ratio and top right the associated P-value, corrected for multiple comparisons using the Šidák correction. Asterisks indicate significant pairwise differences. * = P<0.05, ** = P<0.01, *** = P<0.001.

| | ***H. erato*** | ***H. melpomene*** | ***H. hecale*** | ***A. vanillae*** | ***D. phaetusa*** | ***D. iulia*** |
|---|---|---|---|---|---|---|
| ***H. erato*** | | 0.0050** | 0.0001*** | 0.737 | 0.045* | <0.0001*** |
| ***H. melpomene*** | 3.585** | | 1.000 | 0.225 | 1.000 | 1.000 |
| ***H. hecale*** | 4.439*** | 0.189 | | 0.020* | 0.967 | 1.000 |
| ***Agraulis vanillae*** | 1.721 | -2.391 | -3.213* | | 0.831 | 0.007** |
| ***Dryadula phaetusa*** | 2.964* | -0.889 | -1.274 | 1.591 | | 0.867 |
| ***Dryas iulia*** | 4.688*** | 0.419 | 0.286 | 3.506** | 1.531 | |

**Table S2** Naïve colour preference for each species. Results for each species for generalised linear mixed models including only ID as a random effect. P-values less than 0.05 indicate significant deviation from 50%. * = P<0.05, ** = P<0.01, *** = P<0.001.

| | **z value** | **Pr(>\|z\|)** |
|---|---|---|
| ***H. erato*** | 5.781 | <0.0001*** |
| ***H. melpomene*** | -0.13 | 0.897 |
| ***H. hecale*** | -0.511 | 0.609 |
| ***Agraulis vanillae*** | 4.381 | <0.0001*** |
| ***Dryadula phaetusa*** | 1.163 | 0.245 |
| ***Dryas iulia*** | -0.95 | 0.342 |

### Interspecific variation in associative colour-learning performance

Although all species showed a significant shift in preference towards the trained colour, there were significant differences between species in their fidelity to the trained colour during the first training test (Fig. S6, $\chi^2$=20.536, d.f.=5, P<0.001), with *Dryadula phaetusa* significantly less accurate than *H. erato*, and *Dryas iulia* less accurate than *H. erato* and *Agraulis vanillae nius* species and *Agraulis vanillae* (Fig. S6, Table S3). Overall, however, *Heliconius* individuals were not significantly more accurate than non-*Heliconius* species ($\chi^2$=3.246, d.f.=1, P=0.0716). All species displayed a high fidelity to the trained colour, with the mean proportion of correct attempts ranging from 0.838 in *Dryas iulia* to 0.929 in *H. erato* (Fig. S6).

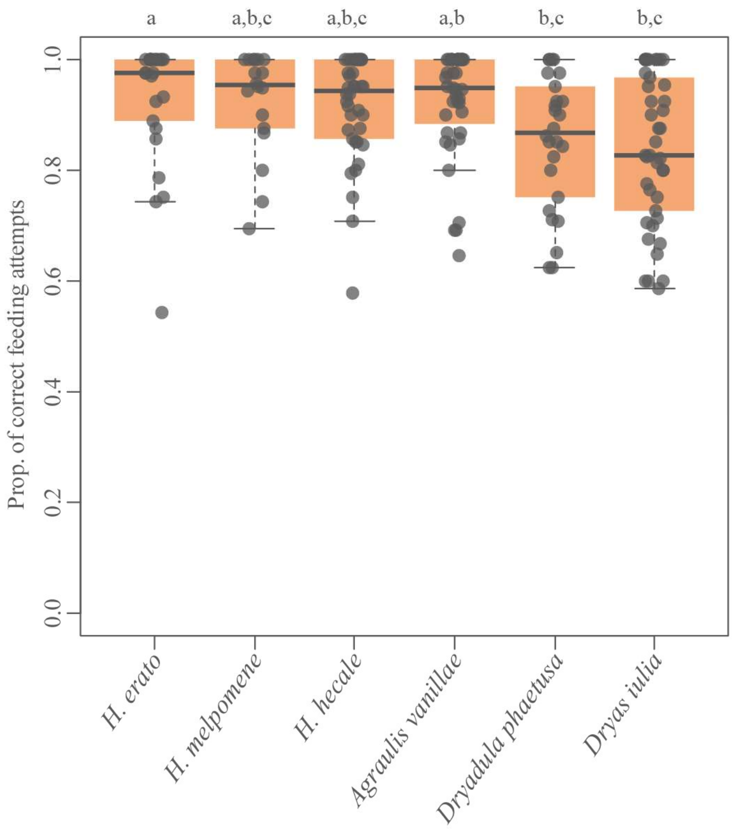


**Fig. S6** 16-hour recall performance in a two-colour preference test after four days of training for three *Heliconius* species (*H. erato* (n=25), *H. melpomene* (n=18) and *H. hecale* (n=38)) and three non-*Heliconius* Heliconiini, *Agraulis vanillae* (n=40), *Dryadula phaetusa* (n=26) and *Dryas iulia* (n=41).

**Table S3** Pairwise comparisons between species in fidelity to the trained colour cue during the first recall test. Bottom left shows z-ratio and top right the associated P-value, corrected for multiple comparisons using the Šidák correction. *=P<0.05.

| | ***H. erato*** | ***H. melpomene*** | ***H. hecale*** | ***A. vanillae*** | ***D. phaetusa*** | ***D. iulia*** |
|---|---|---|---|---|---|---|
| ***H. erato*** | | 0.909 | 0.529 | 0.975 | 0.022* | 0.01* |
| ***H. melpomene*** | 1.028 | | 0.998 | 0.998 | 0.444 | 0.362 |
| ***H. hecale*** | 1.705 | 0.446 | | 0.870 | 0.491 | 0.359 |
| ***Agraulis vanillae*** | 0.755 | -0.441 | -1.128 | | 0.054 | 0.022* |
| ***Dryadula phaetusa*** | 3.124* | 1.833 | 1.762 | 2.820 | | 1.000 |
| ***Dryas iulia*** | 3.364* | 1.967 | 1.971 | 3.132* | 0.025 | |

### Interspecific variation in long-term memory performance

There was significant interspecific variation in performance in the long-term memory test. Both *H. erato* and *H. melpomene* were more accurate than the three non-*Heliconius* species during the long-term memory test (Table S5). However, the long-term memory performance of *H. hecale* was significantly higher than *Dryas iulia*, but not the other non-*Heliconius* species or the other *Heliconius*, suggesting a degree of intermediacy (Table S5). Indeed, uncorrected pairwise comparisons show *H. hecale* as both performing significantly better than both *Dryas iulia* and *Agraulis vanillae*, and worse than the other *Heliconius* (Table S5). The three outgroup Heliconiini, *Dryas iulia*, *Dryadula phaetusa* and *Agraulis vanilla*, did not differ from each other in long-term memory performance (Table S5).

**Table S4** Pairwise comparisons for within species shifts in colour preferences across trials during the long-term memory assay, corrected for multiple comparisons using the Šidák correction. Trained = 16-hour recall after four days of training; LTM = recall after eight days feeding on neutral (white) feeders; LTM2 = recall performance after an additional four days on white feeders.

<table>
<tr><th colspan="2">Species</th><th>Contrast</th><th>z ratio</th><th>P value</th></tr>
<tr><td rowspan="8">Heliconius</td><td rowspan="2">H. erato</td><td>Naïve – Trained</td><td>-10.785</td><td><0.0001***</td></tr>
<tr><td>Trained – LTM</td><td>3.506</td><td>0.0009***</td></tr>
<tr><td rowspan="3">H. melpomene</td><td>Naïve – Trained</td><td>-9.202</td><td><0.0001***</td></tr>
<tr><td>Trained – LTM</td><td>2.558</td><td>0.0313*</td></tr>
<tr><td>LTM-LTM2</td><td>0.230</td><td>0.994</td></tr>
<tr><td rowspan="3">H. hecale</td><td>Naïve – Trained</td><td>-14.035</td><td><0.0001***</td></tr>
<tr><td>Trained – LTM</td><td>6.511</td><td><0.0001***</td></tr>
<tr><td>LTM-LTM2</td><td>1.470</td><td>0.367</td></tr>
<tr><td rowspan="8">Non-Heliconius</td><td rowspan="3">Agraulis vanillae</td><td>Naïve – Trained</td><td>-16.495</td><td><0.0001***</td></tr>
<tr><td>Trained – LTM</td><td>11.446</td><td><0.0001***</td></tr>
<tr><td>LTM-LTM2</td><td>1.070</td><td>0.637</td></tr>
<tr><td rowspan="3">Dryadula phaetusa</td><td>Naïve – Trained</td><td>-9.958</td><td><0.0001***</td></tr>
<tr><td>Trained – LTM</td><td>5.651</td><td><0.0001***</td></tr>
<tr><td>LTM-LTM2</td><td>1.600</td><td>0.294</td></tr>
<tr><td rowspan="2">Dryas iulia</td><td>Naïve – Trained</td><td>-12.303</td><td><0.0001***</td></tr>
<tr><td>Trained – LTM</td><td>8.316</td><td><0.0001***</td></tr>
</table>

**Table S5** Pairwise comparisons between species in recall accuracy after eight days feeding on neural (white) feeders. Top right shows P-values corrected for multiple comparisons using the Šidák correction. Bottom left shows uncorrected P-values. * = P<0.05; ** = P<0.01; *** = P<0.001.

| | ***H. erato*** | ***H. melpomene*** | ***H. hecale*** | ***A. vanillae*** | ***D. phaetusa*** | ***D. iulia*** |
|---|---|---|---|---|---|---|
| ***H. erato*** | | 1.000 | 0.2600 | 0.0002*** | 0.002** | <0.0001*** |
| ***H. melpomene*** | 0.9766 | | 0.425 | 0.0014** | 0.009** | 0.0001*** |
| ***H. hecale*** | 0.0199* | 0.0362* | | 0.258 | 0.662 | 0.018* |
| ***Agraulis vanillae*** | <0.0001*** | 0.0001*** | 0.0197* | | 1.000 | 0.998 |
| ***Dryadula phaetusa*** | 0.0001*** | 0.0006*** | 0.0697 | 0.783 | | 0.989 |
| ***Dryas iulia*** | <0.0001*** | <0.0001*** | 0.0012** | 0.338 | 0.262 | |

**Table S6** Colour preference for each species 13 days after training ended. Results for each species for generalised linear mixed models including only ID as a random effect. P-values less than 0.05 indicate significant deviation from 50% preference between purple and yellow. *** = P<0.001.

| | **z value** | **Pr(>\|z\|)** |
|---|---|---|
| ***H. melpomene*** | 3.745 | 0.00018*** |
| ***H. hecale*** | 1.824 | 0.0682 |
| ***Agraulis vanillae*** | -0.593 | 0.553 |
| ***Dryadula phaetusa*** | -0.82 | 0.412 |

### Control butterflies show no shift in colour preference

For both *Heliconius erato* and *Dryas* iulia, there was no shift in colour preference after four days' exposure to coloured feeders in the "non-learning" control environment (Fig. S7, Table S7), suggesting that they did not form a learned association with either colour.

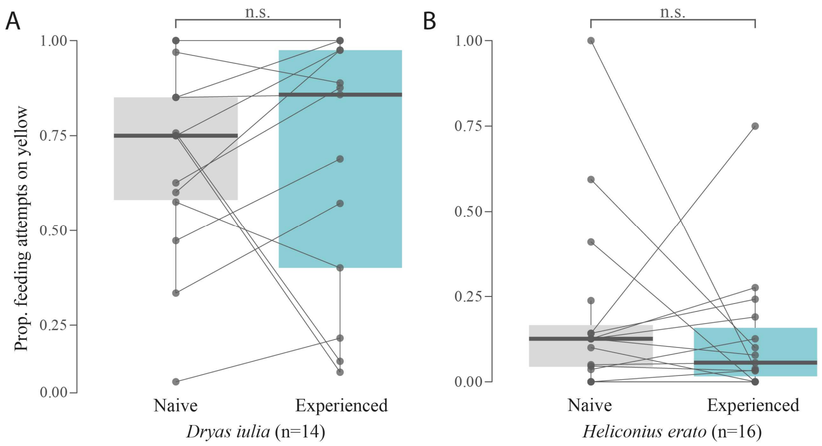


**Fig. S7** Control group butterflies show no shift in colour preference after exposure to coloured feeders in both (*A*) *Dryas iulia* and (*B*) *Heliconius erato*. Control butterflies were kept for four days in an environment where half of the purple feeders were presented with a sugar-pollen reward and half with an aversive quinine solution, likewise for yellow feeders.

**Table S7** Pairwise comparisons between colour preference before and after experience with coloured feeders for Control group *Dryas iulia* and *Heliconius erato*. Correct for multiple comparisons using Tukey's test.

| Species | t ratio | P value |
|---|---|---|
| ***Dryas iulia*** | 0.587 | 0.916 |
| ***H. erato*** | 2.177 | 0.652 |

**Table S8** ANOVA results for a series of GLMMs testing for variation between *Dryas iulia* and *Heliconius erato*, divided into three treatment groups – Day 0, Learning and Control – in several neuroanatomical traits measured in the mushroom body calyx.

| Trait | Factor | $\chi^2$ | Df | Pr(>$\chi^2$) |
|---|---|---|---|---|
| **Calyx synapse density** | **Group** | 33.771 | 2 | <0.0001*** |
| | **Species** | 1.373 | 1 | 0.241 |
| | **Group:Species** | 6.312 | 2 | 0.0426* |
| **Mushroom body calyx volume** | **Group** | 30.170 | 2 | <0.0001*** |
| | **Species** | 771.720 | 1 | <0.0001*** |
| | **Group:Species** | 6.380 | 2 | 0.0413* |
| **Number of calyx synapses** | **Group** | 25.857 | 2 | <0.0001*** |
| | **Species** | 40.056 | 1 | <0.0001*** |
| | **Group:Species** | 10.338 | 2 | 0.0057** |
| **Number of Kenyon cells** | **Group** | 4.790 | 2 | 0.091 |
| | **Species** | 773.640 | 1 | <0.0001*** |
| | **Group:Species** | 0.170 | 2 | 0.918 |

**Table S9** Selected pairwise comparisons in several neuroanatomical traits measured in the mushroom body calyx. Comparisons are made within species between groups, and between species for individuals in the same treatment group. Corrected for multiple comparisons using the Šidák test.

| **Trait** | **Species** | **Contrast** | **t ratio** | **Uncorrected P value** | **Corrected P value** |
|---|---|---|---|---|---|
| **Calyx synapse density** | ***Dryas iulia*** | Day 0 – Learning | 1.265 | 0.209 | 0.505 |
| | | Day 0 – Control | 1.888 | 0.062 | 0.176 |
| | | Learning – Control | 0.932 | 0.354 | 0.730 |
| | ***H. erato*** | Day 0 – Learning | 4.490 | <0.0001*** | <0.0001*** |
| | | Day 0 – Control | 5.916 | <0.0001*** | <0.0001*** |
| | | Learning – Control | 2.163 | 0.033* | 0.097 |
| | ***Dryas iulia* – *H. erato*** | Day 0 | -1.519 | 0.132 | 0.347 |
| | | Learning | 1.387 | 0.169 | 0.426 |
| | | Control | 1.859 | 0.066 | 0.186 |
| **Mushroom body calyx volume** | ***Dryas iulia*** | Day 0 – Learning | -1.971 | 0.0523* | 0.149 |
| | | Day 0 – Control | -1.396 | 0.167 | 0.421 |
| | | Learning – Control | 0.346 | 0.730 | 0.980 |
| | ***H. erato*** | Day 0 – Learning | -5.500 | <0.0001*** | <0.0001*** |
| | | Day 0 – Control | -4.317 | <0.0001*** | <0.0001*** |
| | | Learning – Control | 1.098 | 0.275 | 0.620 |
| | ***Dryas iulia* – *H. erato*** | Day 0 | -12.259 | <0.0001*** | <0.0001*** |
| | | Learning | -19.864 | <0.0001*** | <0.0001*** |
| | | Control | -15.271 | <0.0001*** | <0.0001*** |
| **Number of calyx synapses** | ***Dryas iulia*** | Day 0 – Learning | 0.429 | 0.669 | 0.964 |
| | | Day 0 – Control | 0.973 | 0.334 | 0.704 |
| | | Learning – Control | 0.694 | 0.490 | 0.867 |
| | ***H. erato*** | Day 0 – Learning | 3.519 | 0.0007*** | 0.0022** |
| | | Day 0 – Control | 5.928 | <0.0001*** | <0.0001*** |
| | | Learning – Control | 2.700 | 0.0064** | 0.0254* |
| | ***Dryas iulia* – *H. erato*** | Day 0 | -5.783 | <0.0001*** | <0.0001*** |
| | | Learning | -3.872 | 0.0002 | 0.0007*** |
| | | Control | -1.399 | 0.166 | 0.420 |
| **Number of Kenyon cells** | ***Dryas iulia*** | Day 0 – Learning | -1.436 | 0.155 | 0.397 |
| | | Day 0 – Control | -1.419 | 0.160 | 0.408 |
| | | Learning – Control | -0.202 | 0.841 | 0.996 |
| | ***H. erato*** | Day 0 – Learning | -0.918 | 0.362 | 0.740 |
| | | Day 0 – Control | -1.540 | 0.123 | 0.337 |
| | | Learning – Control | -0.674 | 0.503 | 0.877 |
| | ***Dryas iulia* – *H. erato*** | Day 0 | -14.799 | <0.0001*** | <0.0001*** |
| | | Learning | -17.964 | <0.0001*** | <0.0001*** |
| | | Control | -15.235 | <0.0001*** | <0.0001*** |

**Table S10** Pairwise comparisons for changes in elevation in the scaling relationship between Kenyon cell number and synapse number in the calyx, and Kenyon cell number and calyx volume.

| Regression | Species | Contrast | Test stat | Uncorrected P value | Corrected P value |
|---|---|---|---|---|---|
| **Number of calyx synapses ~ Number of Kenyon cells** | ***Dryas iulia*** | Day 0 – Learning | 2.384 | 0.123 | 0.859 |
| | | Day 0 – Control | 5.111 | 0.0237* | 0.303 |
| | | Learning – Control | 0.568 | 0.451 | 0.999 |
| | ***H. erato*** | Day 0 – Learning | 7.431 | 0.0064** | 0.092 |
| | | Day 0 – Control | 24.969 | <0.0001*** | <0.0001*** |
| | | Learning – Control | 5.268 | 0.0217* | 0.327 |
| | ***Dryas iulia* – *H. erato*** | Day 0 | 4.866 | 0.0274* | 0.341 |
| | | Learning | 26.805 | <0.0001*** | <0.0001*** |
| | | Control | 18.126 | <0.0001*** | 0.0003*** |
| **Calyx volume ~ Number of Kenyon cells** | ***Dryas iulia*** | Day 0 – Learning | 0.526 | 0.468 | 0.999 |
| | | Day 0 – Control | 0.0003 | 0.986 | 1.000 |
| | | Learning – Control | 0.0300 | 0.862 | 1.000 |
| | ***H. erato*** | Day 0 – Learning | 9.727 | 0.0018** | 0.0269* |
| | | Day 0 – Control | 5.489 | 0.0191* | 0.252 |
| | | Learning – Control | 2.557 | 0.110 | 0.825 |
| | ***Dryas iulia* – *H. erato*** | Day 0 | 0.625 | 0.429 | 0.999 |
| | | Learning | 0.00288 | 0.957 | 1.000 |
| | | Control | 0.618 | 0.432 | 1.000 |

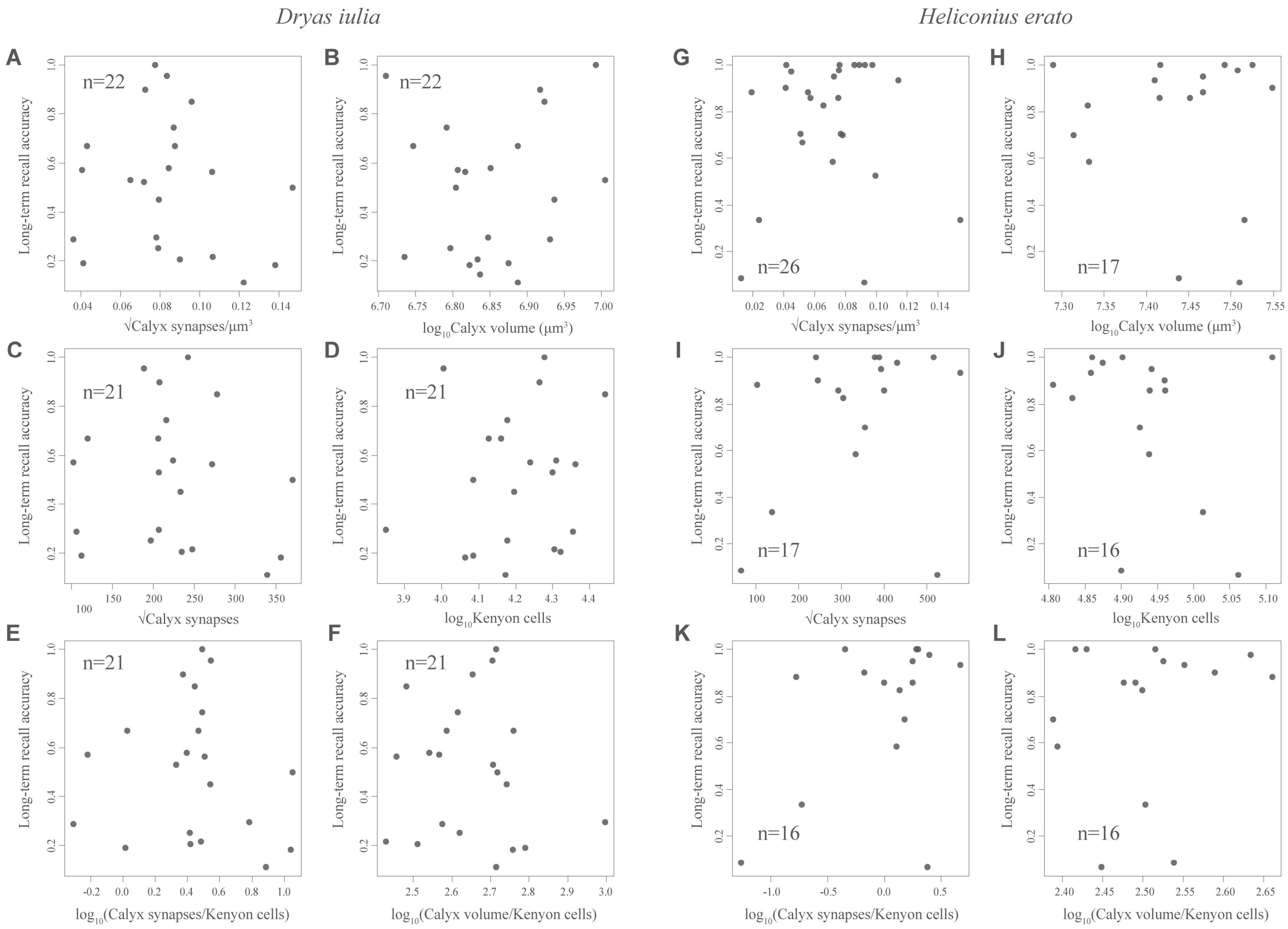


**Fig. S8** Individual performance in the long-term recall test (after eight days fed on neutral (white) feeders) is not correlated with any of the traits measured in the mushroom body calyx for either (*A*)-(*F*) *Dryas iulia* or (*G*)-(*L*) *Heliconius erato*.

**Table S10** ANOVA results for a series of GLMMs testing for relationships between neuroanatomical measurements in the mushroom body calyx and performance in the initial and long-term recall tests in *Dryas iulia* and *Heliconius erato*.

| Trait | Test | Species | $\chi^2$ | Df | Pr(> $\chi^2$) |
|---|---|---|---|---|---|
| **Synapse density** | **Initial recall** | ***Dryas iulia*** | 1.4775 | 1 | 0.224 |
| | | ***H. erato*** | 5.473 | 1 | 0.0193* |
| | **Long-term recall** | ***Dryas iulia*** | 0.4562 | 1 | 0.499 |
| | | ***H. erato*** | 0.1482 | 1 | 0.700 |
| **Mushroom body calyx volume** | **Initial recall** | ***Dryas iulia*** | 7.6747 | 1 | 0.0056** |
| | | ***H. erato*** | 0.676 | 1 | 0.411 |
| | **Long-term recall** | ***Dryas iulia*** | 0.0368 | 1 | 0.848 |
| | | ***H. erato*** | 0.099 | 1 | 0.753 |
| **Number of calyx synapses** | **Initial recall** | ***Dryas iulia*** | 0.3431 | 1 | 0.558 |
| | | ***H. erato*** | 9.199 | 1 | 0.0024** |
| | **Long-term recall** | ***Dryas iulia*** | 0.3575 | 1 | 0.550 |
| | | ***H. erato*** | 1.585 | 1 | 0.208 |
| **Number of Kenyon cells** | **Initial recall** | ***Dryas iulia*** | 0.0068 | 1 | 0.934 |
| | | ***H. erato*** | 1.585 | 1 | 0.208 |
| | **Long-term recall** | ***Dryas iulia*** | 0.4704 | 1 | 0.493 |
| | | ***H. erato*** | 1.5167 | 1 | 0.218 |
| **Ratio of synapses to Kenyon cells** | **Initial recall** | ***Dryas iulia*** | 0.576 | 1 | 0.448 |
| | | ***H. erato*** | 12.852 | 1 | 0.00034*** |
| | **Long-term recall** | ***Dryas iulia*** | 0.2323 | 1 | 0.630 |
| | | ***H. erato*** | 3.2646 | 1 | 0.0708 |
| **Ratio of calyx volume to Kenyon cells** | **Initial recall** | ***Dryas iulia*** | 1.8266 | 1 | 0.177 |
| | | ***H. erato*** | 0.2405 | 1 | 0.624 |
| | **Long-term recall** | ***Dryas iulia*** | 0.3875 | 1 | 0.534 |
| | | ***H. erato*** | 0.3793 | 1 | 0.538 |

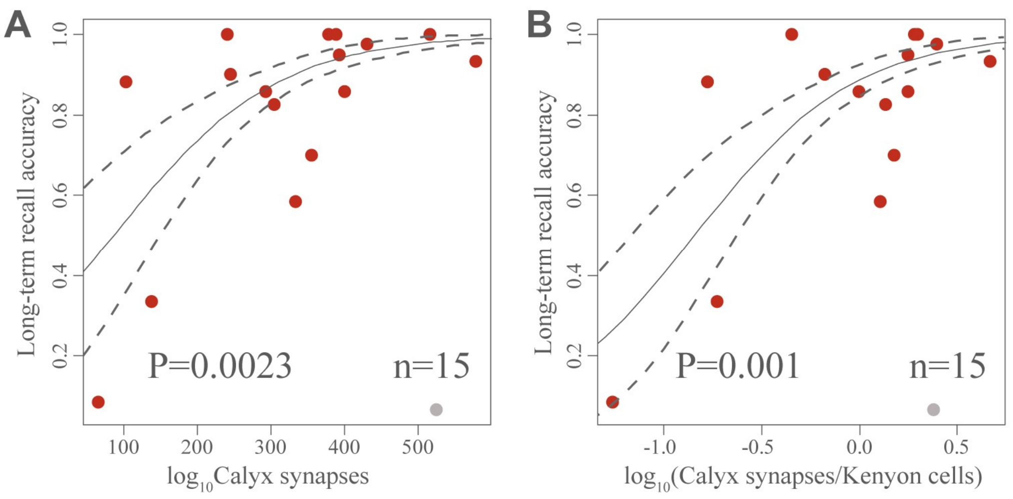


**Fig. S9** Long-term recall (8 days) accuracy and synapse number in *Heliconius erato*. When a single outlier individual (grey) is removed, (*A*) the total synapse number in the calyx ($\chi^2$=9.263, d.f.=1, P=0.002) and (*B*) the ratio of synapses to Kenyon cells ($\chi^2$=10.918, d.f.=1, P=0.001) show a significant positive relationship with performance in the long-term recall test. The outlier individual exhibited highly unusual behaviour, making 16 out of 16 correct feeding attempts in the initial recall test, but only 2 out of 28 correct attempts in the long-term recall test after eight days with neutral stimuli. Regression lines, with standard errors, are from GLMM analysis.
